# Nanopore sequencing undergoes catastrophic sequence failure at inverted duplicated DNA sequences

**DOI:** 10.1101/852665

**Authors:** Pieter Spealman, Jaden Burrell, David Gresham

## Abstract

Inverted duplicated sequences are a common feature of structural variants (SVs) and copy number variants (CNVs). Analysis of CNVs containing inverted duplicated sequences using nanopore sequencing identified recurrent aberrant behavior characterized by incorrect and low confidence base calls that result from a systematic elevation in the current recorded by the sequencing pore. The coincidence of inverted duplicated sequences with catastrophic sequence failure suggests that secondary DNA structures may impair transit through the nanopore.

---

Advances in DNA sequencing have led to a greater understanding of the role structural variants play in disease [1, 2, 3] and evolution [4]. However, high throughput short read DNA sequencing faces intrinsic challenges in identifying complex structural variants. A variety of computational methods use sequence read depth, split, and discordant reads to identify structural variants. However, these methods are uninformative in low complexity and repetitive regions of the genome [5, 2, 6]. Long-read DNA sequencing can potentially overcome these technical limitations by generating reads of 10-100 kilobases (kb) length that enable unambiguous resolution of contiguous DNA sequences [7, 8, 9]. Nanopore sequencing, developed by Oxford-Nanopore Technologies (ONT), comprises a sequencing pore that translocates DNA or RNA across a membrane and sensors that measure changes in ionic current during the translocation.The electrical signals are subsequently converted into base-called sequences using a machine-learning base-caller [8, 10]. The very long sequence reads that can be generated using this technology has proven invaluable for identifying many types of structural variants (SVs), such as translocations, tandem duplications, and simple inversions which are difficult to identify using short-read sequencing alone [11, 12].

Inverted duplications are a class of copy-number variants (CNVs) that feature two duplicated regions oriented in opposite directions with a junction between them. Inverted repeated DNA sequences occur frequently in many genomes. They have been identified as playing a role in adaptation and evolution [13, 14] and have been associated with human pathologies and diseases [15, 16].

Here, we show that genomic regions containing large inverted duplications generate nanopore reads with pronounced base-calling errors and low sequence quality (i.e. phred scores). We propose that this high error rate is caused by biophysical interference induced upon the translocating DNA by the formation of large-scale secondary structures at inverted repeat sequences. As the specific secondary structures formed by these sequences are likely to depend on the length and nucleotide composition of the inverted repeat, it may be impossible to learn a generalizable signal for these sequences, presenting an intractable problem for nanopore sequencing.

## Results

SVs and CNVs result in novel sequences that are often diagnostic of the formation mechanism [3]. Inverted repeat sequences are CNVs that feature repeated sequences oriented in the opposite directions with a junction region separating them (**Figure 1b**). We recently identified numerous CNVs containing inverted repeat sequences (**Figure 1a**) in experimentally evolved strains of *Saccharomyces cerevisiae* using short-read Illumina sequencing [14]. These CNVs are likely formed by origin dependent inverted repeat amplification (ODIRA), a DNA replication-based mechanism of CNV formation [13, 17]. ODIRA-generated CNVs are characterized by triplicated sequence, of hundreds to thousands of bases, with the internal repeat inverted relative to the other two copies. As the junction between repeated sequences results in a unique sequence it enables their identification using short-read sequencing. The structure of ODIRA breakpoints can be verified using orthogonal methods such as restriction digest analysis and Southern blotting [17].

**Figure 1.**
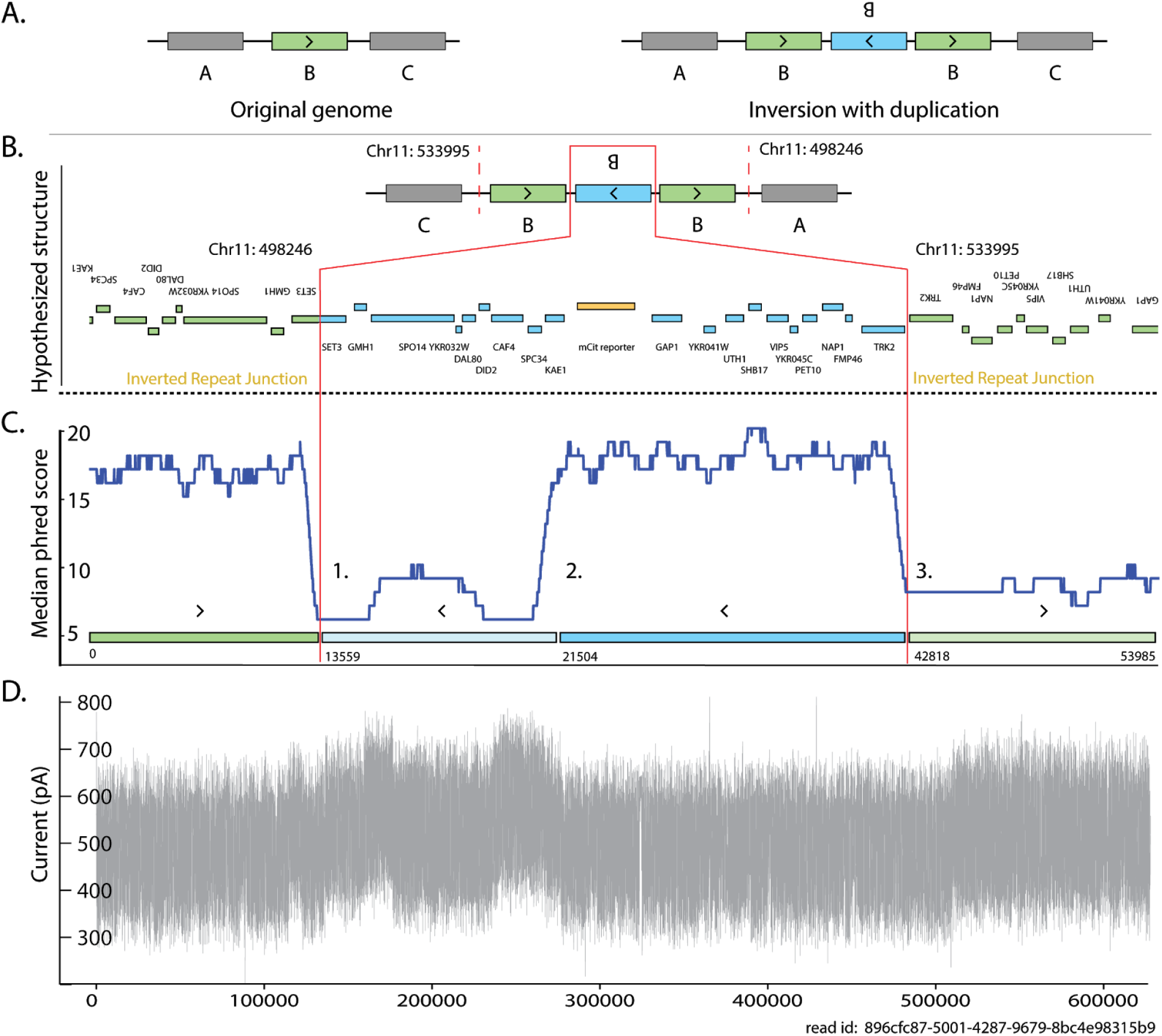
Schematic of a long read sequence generated using nanopore sequencing that spans two inverted repeat junctions. This example shows a single read spanning a known CNV containing a triplication that comprises two inverted duplications **(A)**. The hypothesized structure based on the alignment of Illumina sequencing reads to the reference genome [14] **(B)**. The predicted inverted repeat junctions [14] are denoted with two vertical red lines. A rolling 1000 base window median phred score is calculated for the read **(C)**. The low-phred scoring regions begin at the predicted junction (**Points 1 and 3**) and recovers after a distance similar to the length of the preceding normal quality region (**2**). An analysis of the raw current generated by the nanopore during the translocation of the DNA molecule shows a substantial uplift in current co-occurs around the low-phred scoring regions and the junction between inverted duplicated sequence **(D)**.

We sought to use MinION long read sequencing to verify potential ODIRA CNVs. However, we found that a majority of reads (80-100%) that span the predicted inverted repeat junction exhibited low-phred scores and high-rates of mismatches with the reference genome (**Supplemental Figure 1**). Furthermore, these aberrant reads clustered at the predicted junctions (**Supplemental Figure 2**). To ensure that sequencing failure at inverted repeat junctions was not the product of library construction we verified that there was no substantial variance in sequencing quality (as defined by the phred score) between samples with and without inverted repeat junctions (**Supplemental Figure 3**) Analysis of sequence reads that span inverted duplication junctions showed that the drop in sequencing quality occurred close to junction sequence identified using short read sequencing (**Figure 1c**). In strains that do not contain inverted repeat sequence at the site there is no loss of sequence quality at the corresponding site and the sequence can be unambiguously determined (**Supplemental Figure 4**).

We hypothesized that the low sequence quality at inverted repeat junctions may reflect a degradation in the quality of the nanopore signal. An analysis of the raw current output from the sequencing pore revealed that the decrease in phred score corresponds with an “uplift” in current [18] (**Figure 1d**, **Supplemental Figure 5**). We find that the length of the uplifted regions and corresponding reduced phred score have a high degree of correlation the length of sequence that precedes the inversion junction (**Figure 2**), which is otherwise of typical quality. This high degree of correlation between the lengths of high quality and low quality bases within a single read was only observed for structural variants that have inverted duplicated sequence (**Supplemental Table 3, 4**). We attempted to use a variety of base callers on reads with uplifted regions, but found that no base caller can accurately perform base-calling within these regions. Because nanopore sequencing relies on machine learning to perform base-calling (eg. Albacore and Guppy [10]), the accuracy of base-calling is dependent on how similar the input is to the training data. Because the current in the uplifted region exceeds the range of signals used to train the base caller, the algorithm is unable to perform well, resulting in miscalls, skipped bases, and low phred scores.

**Figure 2.**
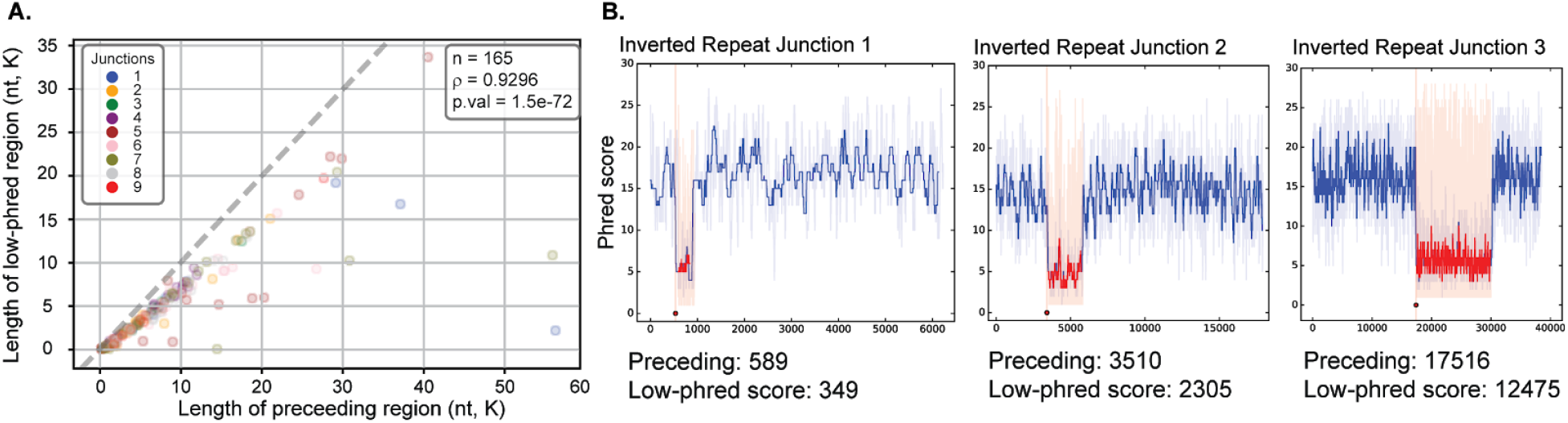
The length of the preceding sequenced region determines the length of the low phred scoring region. For each predicted inverted repeat junction (**Supplemental Table 3**) we compared the length of the low-phred score region with the length of the preceding region that has a typical phred score for all independent sequence reads (see Methods). We find that the lengths of the high quality and subsequent low quality region for each unique read are significantly positively correlated (Spearman *rho* = 0.93, p-value 1.5e-72). For all sequence reads, the preceding region is of slightly greater length than the low-phred scoring regions (the dashed line is the line of identity). This is consistent with the low-scoring region consisting of both miscalled bases and missed bases, potentially due to loss of resolution between bases.

## Discussion

We have identified a phenomenon in which inverted repeat sequences result in a catastrophic decrease in the sequence quality of nanopore long reads, confounding the ability to define this class of SVs using MinION sequencing. We show that the loss of sequence quality is a result of a systematic uplift in the detected current that is the raw signal used for base calling and that the duration of the aberrant signal is a function of the length of the preceding sequence that is of otherwise normal quality.

Because current is a property of translocation rate [19] we propose that the uplift in current is consistent with altered rates of DNA translocation through the pore. Translocation rates could be altered by the intramolecular base-pairing of the complementary sequences on either side of the junction acting as ratchet-like mechanism on the DNA strand or as physical interference generated by interactions between the secondary structures and the nanopore (**Figure 3**). This phenomenon is similar to the observed current uplift in ONT’s now discontinued 2D chemistry which used small oligo linkers to join double-stranded DNA into large self-complementary single strands [18]. Further support for this model is the proximity of the inverted repeat junction to the low-phred scoring regions; and the high correlation between the region lengths before and after the junction.

**Figure 3:**
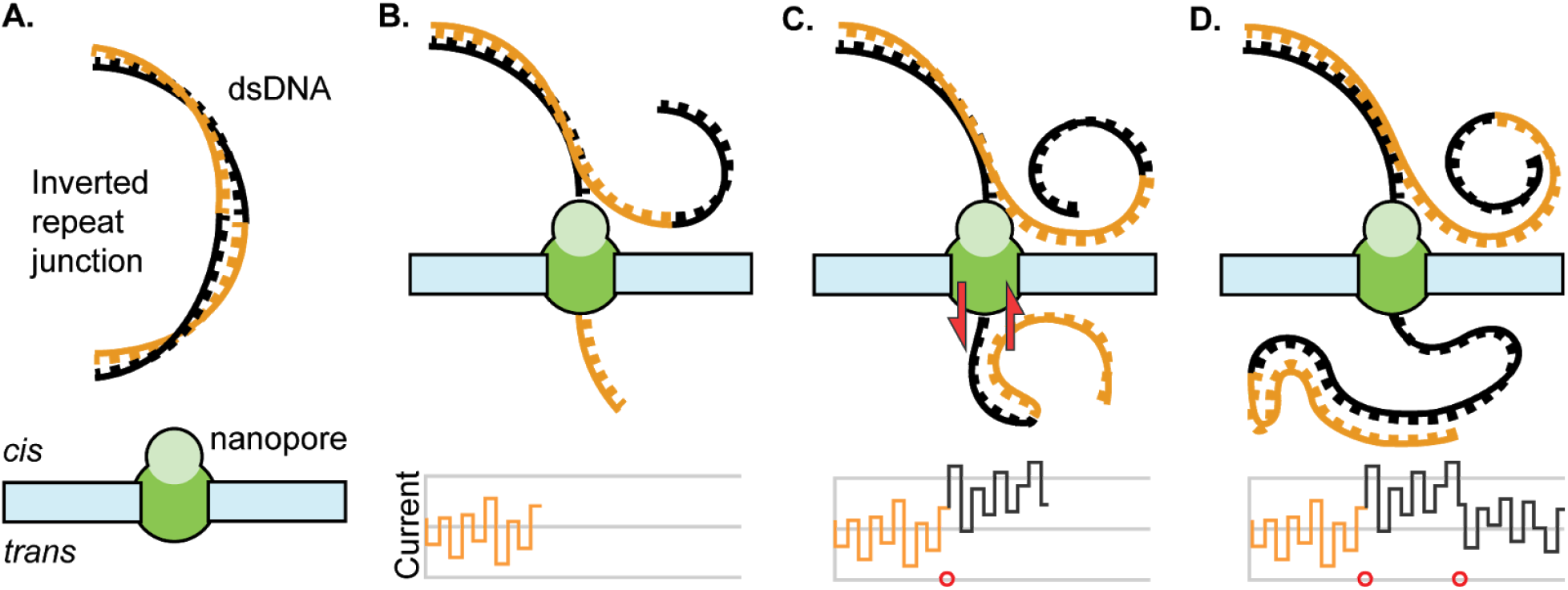
Proposed model for DNA secondary structure interference with sequencing pore. Double stranded DNA containing an inverted duplication (orange and black strands) is introduced to a sequencing pore **(A)**. As the strand translocates through the pore a current is recorded **(B)**. Once the junction has passed to the *trans* side of the nanopore the capacity for the inverted duplication to form secondary structures increases dramatically **(C)**. These secondary structures may increase the pull on the translocating strand or otherwise biophysically interfere with sequencing (red arrows), thus increasing the current generated (first red circle). Once the translocating strand is past the inverted duplication region the interference decreases and the current returns to standard ranges (second red circle) **(D)**. It is also possible that base complementarity on the *cis*- side contributes to impaired transit through the pore, although the activity of the helicase may negate the effect these secondary structures have on the sequencing pore.

Our biophysical model for this aberrant behavior suggests that the frequency and magnitude of sequencing failures will be dependant on parameters that drive secondary structure formation, such as temperature, salt concentrations, fragment size, and the capacity of any given sequence to self-anneal, all of which will make the generation of a generalizable machine learning model difficult. Notably, because inverted duplications lack specific sequences, solutions developed that target specific sequences [18] are infeasible. One possible strategy would be to use a biophysical or chemical approach to decrease the rate of secondary structure formation, for example, by using asymmetric salt concentrations to decrease the free-energy of base-pairing [20]. However, the buffers provided by the commercial supplier are of unknown composition making investigation of this possibility currently infeasible.

Finally, we note that we identified this aberrant behavior by virtue of targeted analysis of predicted inverted duplicated sequence. As most analysis tools exclude low quality base calls, we expect that inverted duplicated sequences in genomes of unknown structure may be missed using current approaches.

## Methods and Materials

### Library preparation

All yeast strains were grown to greater than 1 x 10^7^ cells/mL in 30mL minimal media [14] or YPD (**Supplemental Table 1**). Genomic DNA (gDNA) from each strain was extracted using Qiagen 20/G Genomic tips from ~1.5×10^9^ cells using the manufacturer’s protocol.

All gDNA was barcoded using Oxford-Nanopore’s native barcoding genomic DNA kit (EXP-NBD104), adapters were added using the ligation sequencing kit (SQK-LSK109). The manufacturer’s protocol (versions NBE_9065_v109_revB_23May2018 and NBE_9065_V109_revP_14Aug2019) was followed with the following exceptions: Incubation times for enzymatic repair step were increased to 15min. All Agencourt AMPure XP beads were incubated for 30min at 37°C before elution. Adapter ligations time was increased to 10min. Multiplexed libraries were loaded on MinION flowcells (FLO-MIN106D R9) and run on a MinION sequencer (MIN-101B).

### Base-calling and Alignment

Base-calling was performed using Albacore v2.3.4, Guppy v2.3.5, and Guppy v3.0.3. Low-phred scores associated with junctions of inverted duplicated sequences were observed using each software. All subsequent analysis was performed using Guppy v3.0.3 with the ‘--fast5_out’ option enabled to save the associated squiggle data for each read. The raw electrical signal (e.g. squiggle) was extracted using SquiggleKit [21]. Demultiplexing was performed using Epi2Me with default settings.

All alignments were made using minimap2 with the ‘-ax map-ont’ option [22].

### Low-phred score analysis and reflection point identification

We developed a bioinformatics pipeline, Mugio (https://github.com/pspealman/mugio), that uses fastq and bam files to visualize and quantify reads with low-phred scores and identify potential inverted repeat sequence junctions. Because of the prevalence of reads with low phred scoring regions (**Supplemental Table 2**) candidate junctions are required to meet several criteria. First, reads are scanned to identify reads that map to the same chromosome in different orientations. The reference coordinates that mark the boundaries of split-reads are stored as pairs with overlapping or conflicting pairs resolved using a greedy algorithm. Finally, candidate junctions must have a significant asymmetry in read-depth upstream and downstream of the junction.

To measure the read length pre-junction and post-junction of low phred scoring regions, we first filtered reads that did not overlap a predicted inverted repeat junction. Each remaining read was scanned to identify all subregions (minimum 10 bases in length) that have a median phred score lower than five standard deviations from the global median. Subregions are expanded until the median phred-score in a 10 base window increases above the global median at which point the subregion is stored. Unbounded regions, which never recover from the low-phred score region before the read ends, were not included in length measurements. Pre-junction region lengths were measured from the beginning of the read to the beginning of the low-scoring region.

### Data Availability

Sequence files (Fastq) for all strains referenced in this work are available online at: https://osf.io/qmdwp/. All computer code is available in the github repository: https://github.com/pspealman/mugio

## Acknowledgements

This work was supported by an F32 (1F32GM131573) toPS and funding from the NIH (2R01GM107466) and NSF (MCB1818234) to DG.

## Supplementary Figures

**Supplemental Figure 1.**
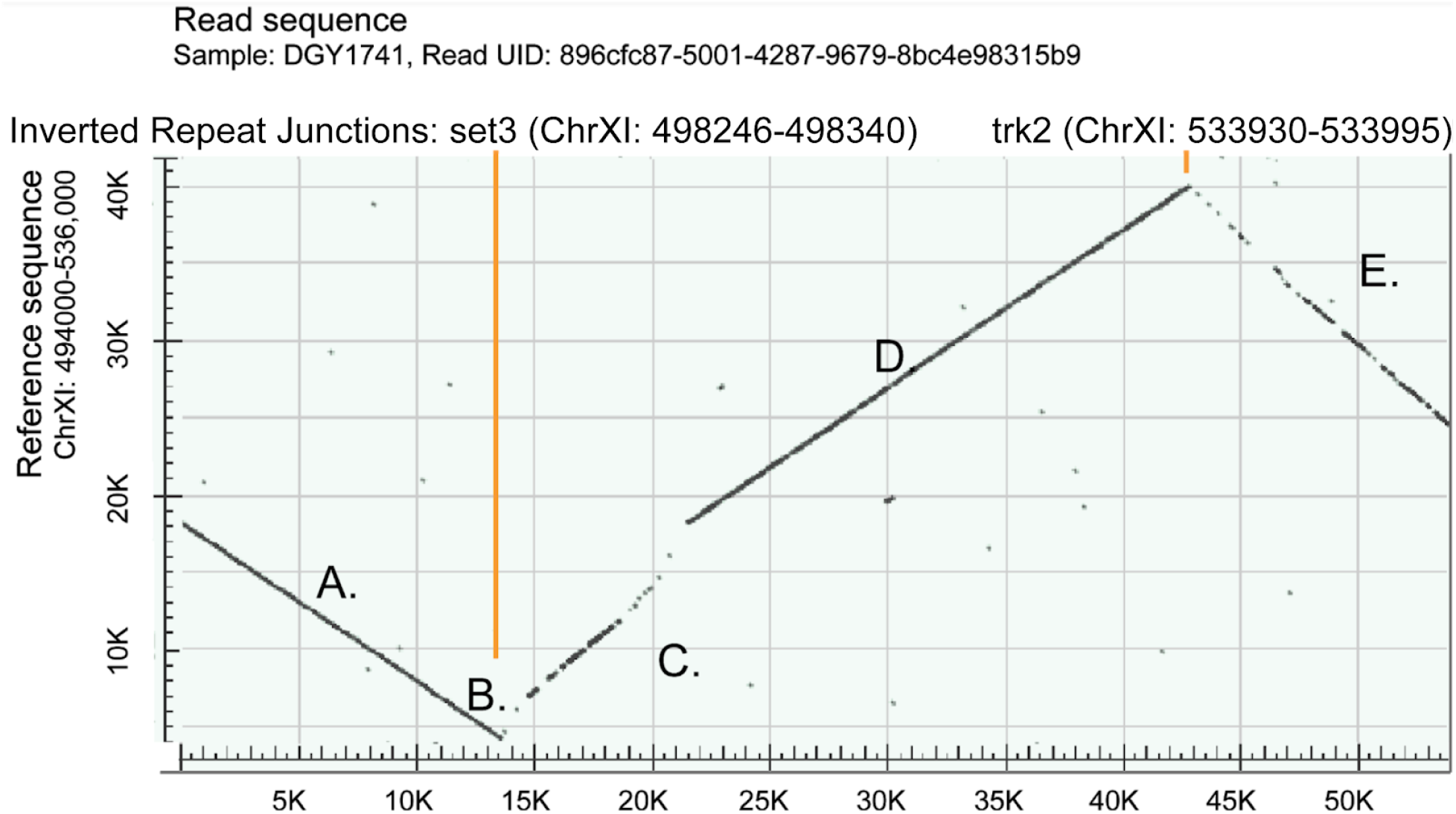
Dot-plot comparison of a read spanning two predicted inverted repeat junctions relative to the reference sequence. The horizontal axis represents the sequence of a single long read (UID: 89ccfc87-5001-4287-9679-8bc4e98315b9) that spans two inverted repeat junctions (orange lines). This is the same read shown in **Figure 1**. The reference sequence is from ChrXI: 494,000 - 536,000. Significant sequence matches between the two are represented by a black dot. The initial solid down-slopping segment represents a close alignment between the read and the negative strand of the reference (**A**). At the inverted repeat junction the direction of the slope changes denoting the inversion (**B**). Note that the length corresponding to the preceding sequence is poorly aligned and of shorter length than the reference (**C**). Once the sequence is past the duplicated region that is contained within the DNA molecule the alignment improves (**D**) until it encounters the second inverted repeat junction when the alignment again degrades (**E**). The read ends before the sequence again recovers.

**Supplemental Figure 2.**
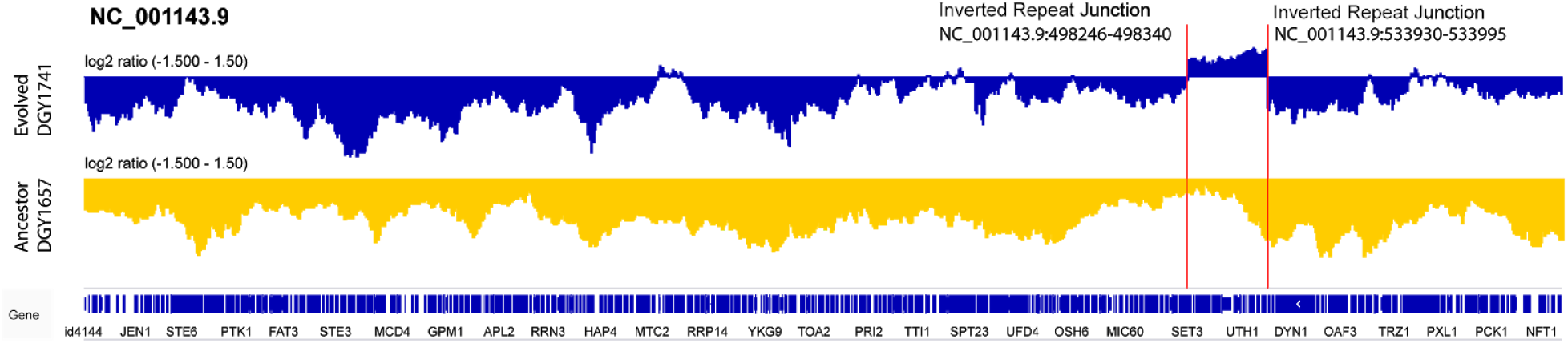
Clustering of reads containing a contiguous span of significant low phred scoring regions. The distribution of reads with significant low phred-scoring regions (see Methods) at the same locus in two yeast strains. For each strain we calculated the log2 transformed ratio of low phred scoring reads (reads with a minimum of length of 50 contiguous nucleotides with a phred score greater than 3 standard deviations from the global median) to total reads, using Deeptools (1). The bottom track shows the read distribution for the wildtype strain which lacks inverted repeat junctions. The top track shows the read distribution for a strain that has inverted repeat junctions (red lines). The pronounced peak of reads with significant low phred-scoring regions clustered around the two junctions in the evolved strain this is entirely absent in the wildtype strain. 1. Ramírez, Fidel, Devon P. Ryan, Björn Grüning, Vivek Bhardwaj, Fabian Kilpert, Andreas S. Richter, Steffen Heyne, Friederike Dündar, and Thomas Manke. deepTools2: A next Generation Web Server for Deep-Sequencing Data Analysis. Nucleic Acids Research (2016). doi: 10.1093/nar/gkw257

**Supplemental Figure 3.**
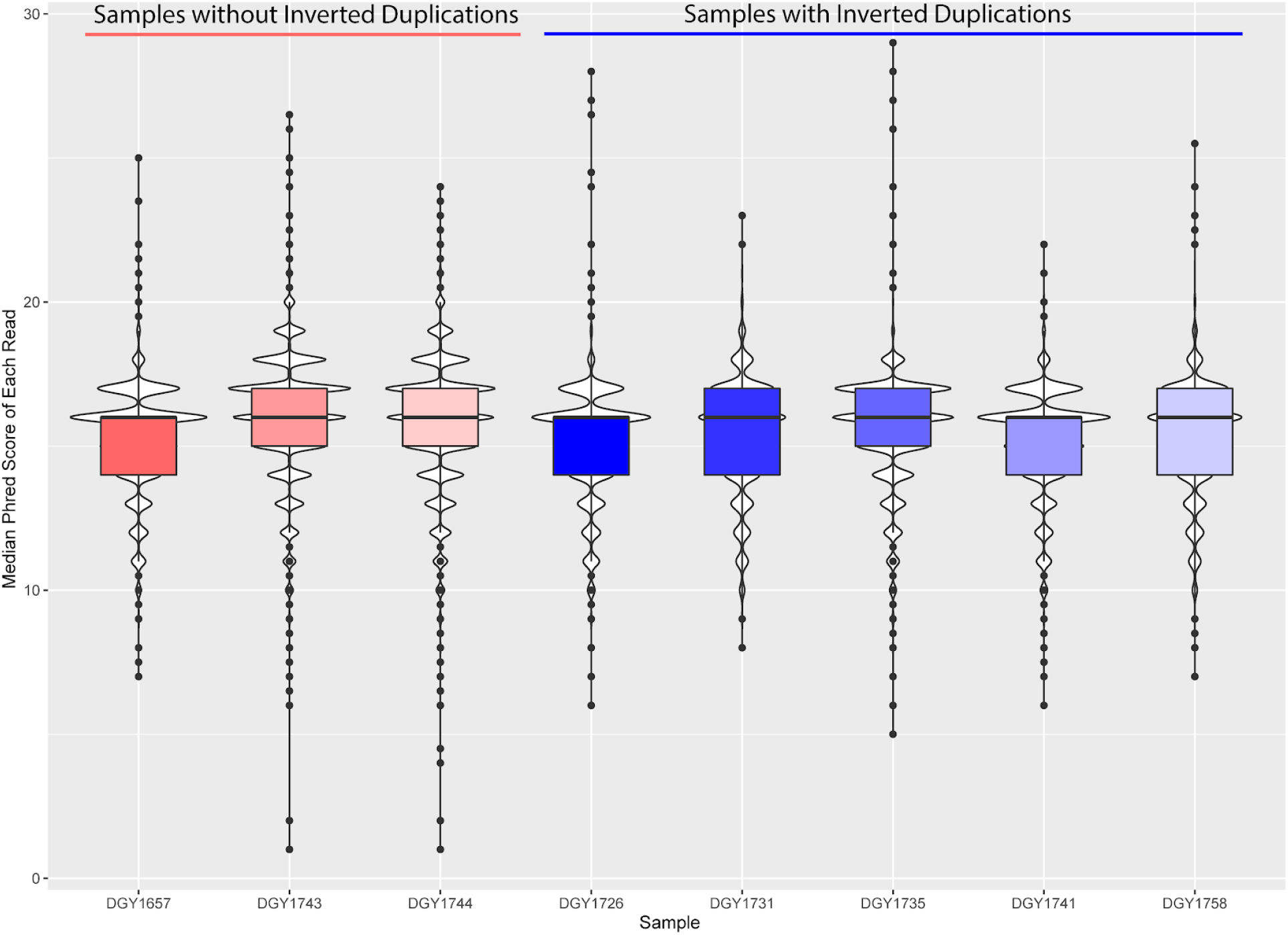
Sequence quality is not globally compromised in genomes containing inverted repeat sequences. The distribution of phred scores of all reads for each sample. The mean of every distribution is within 1.05-fold of the mean of a wildtype strain that does not contain any SVs. There is no substantial difference between the samples without predicted inverted repeats (red) and those with (blue).

**Supplemental Figure 4.**
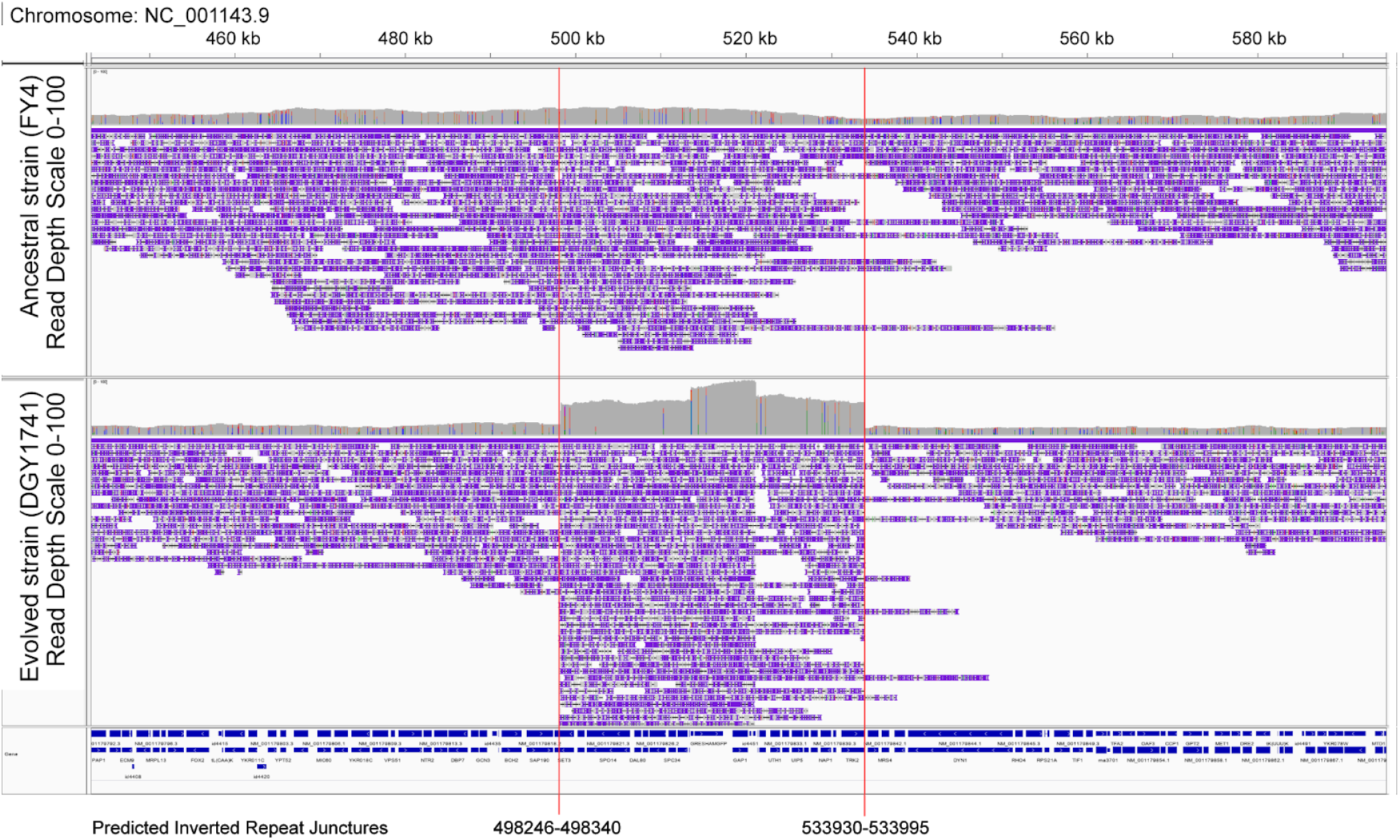
Frequency of low-phred scoring reads at a site with inverted repeat junctions. A comparison of aligned reads with significant low-phred scoring regions (reads with a minimum of length of 50 contiguous nucleotides with a phred score greater than 3 standard deviations from the global median) between the wildtype strain (top track) and a strain with predicted inverted repeat junctions (bottom track). Note the substantially higher abundance of low-scoring reads in thestrain with predicted inverted repeat junctions.

**Supplemental Figure 5.**
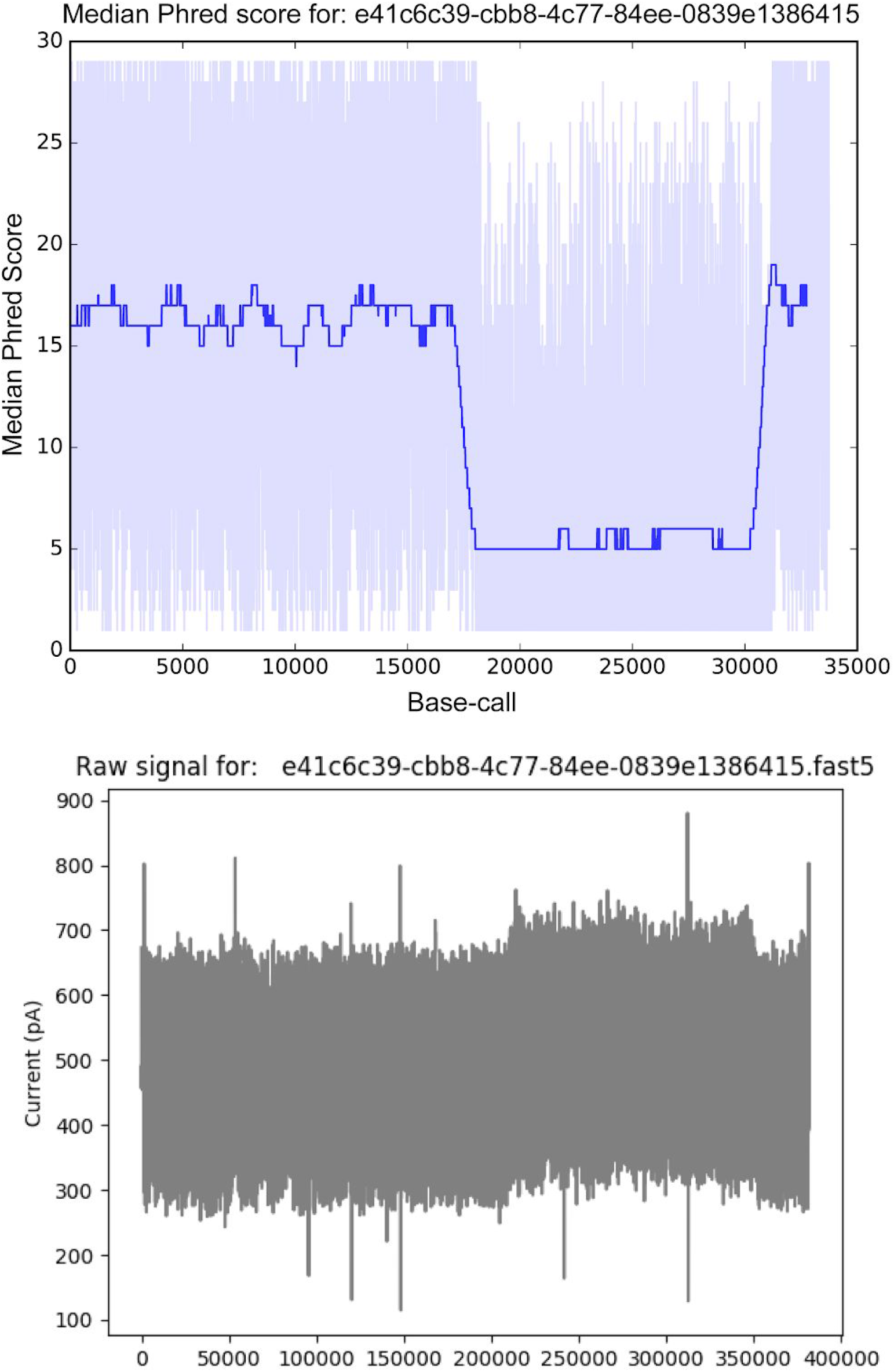
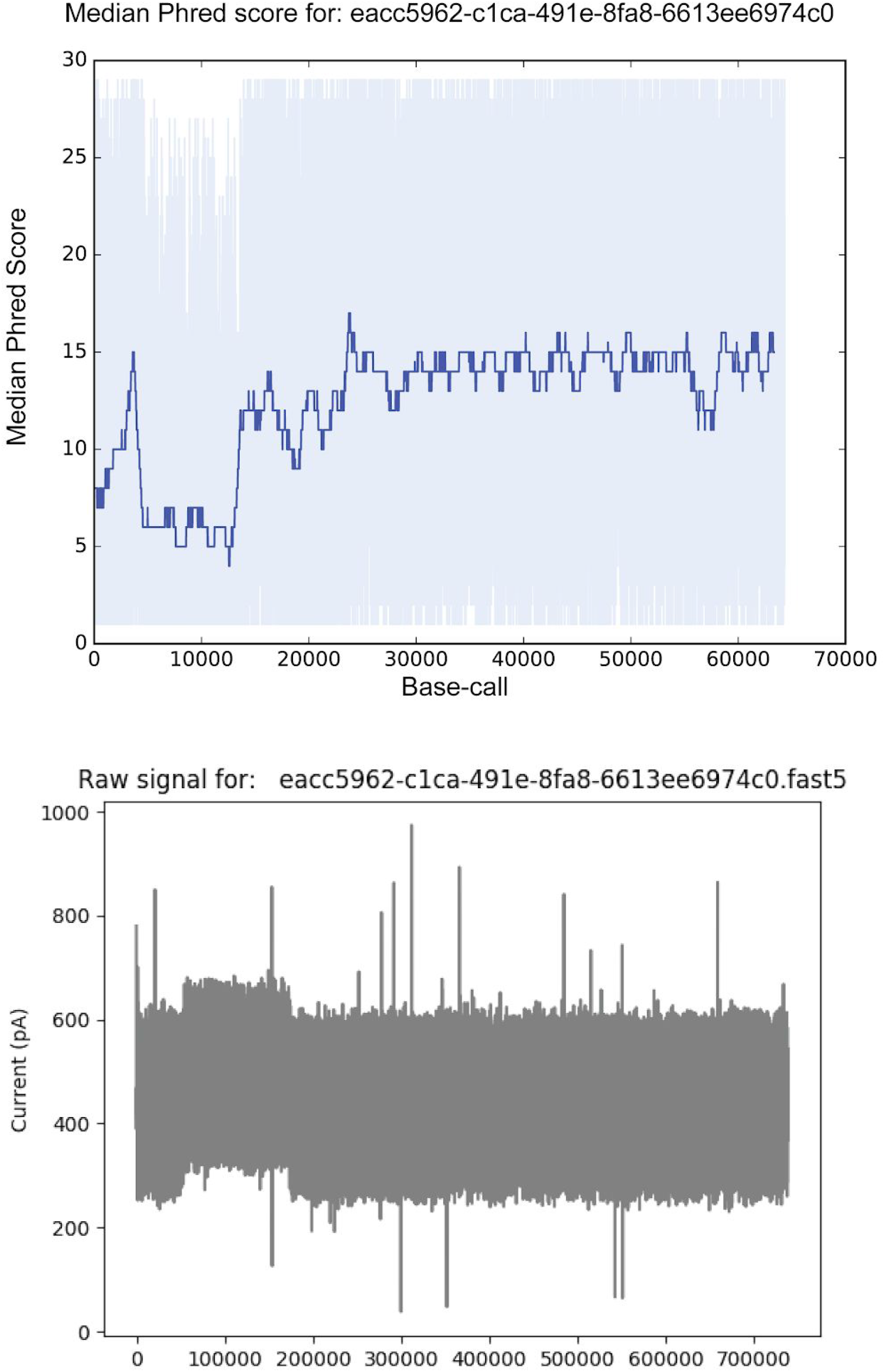
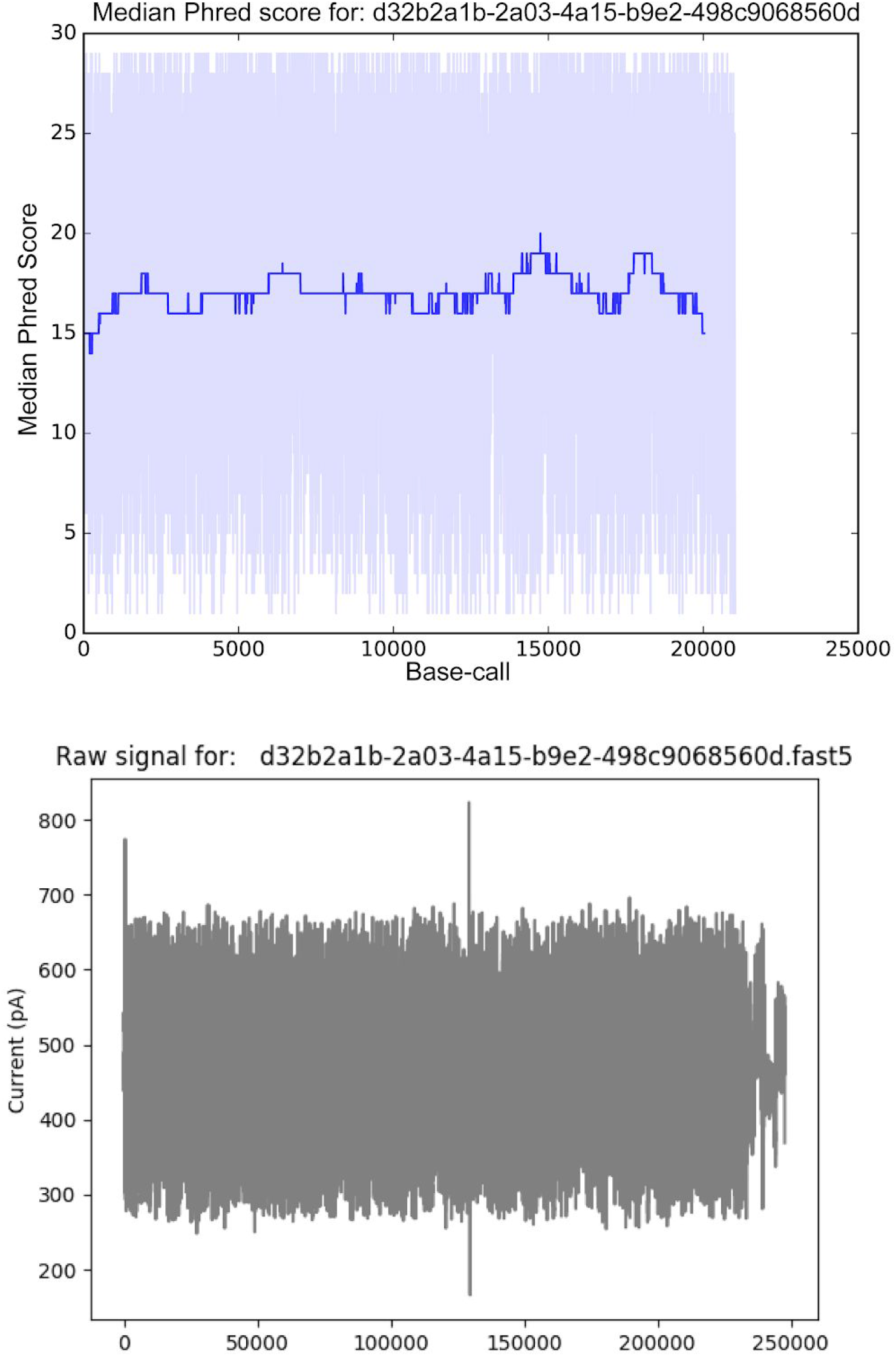
Additional paired windowed phred-score and nanopore raw squiggle results. The first two examples have inverted duplicate repeat junctions, resulting in coincident decreases in phred score and an aberrant squiggle readout. The third example is a read from a region without an inverted repeat junction that lacks both of these properties.

